# Beyond Traditional Methods: Innovative Integration of LISS IV and Sentinel 2A Imagery for Unparalleled Insight into Himalayan Ibex Habitat Suitability

**DOI:** 10.1101/2023.07.18.549476

**Authors:** Ritam Dutta, Bheem Dutt Joshi, Vineet Kumar, Amira Sharief, Saurav Bhattcharjee, Rajappa Babu, Mukesh Thakur, Lalit Kumar Sharma

**Affiliations:** Zooological Survey of India, Prani Vigyan Bhawan, Kolkata, West Bengal 700053; University of Madras, Navalar Nagar, Chepauk, Triplicane, Chennai, Tamil Nadu 600005; Southern Regional Centre, Zoological Survey of India, Chennai, Tamil Nadu 600028

**Keywords:** Image classification, Integration image, Ensemble species distribution model, Himalayan Ibex.

## Abstract

Despite advancements in remote sensing, satellite imagery is underutilized in conservation research. Multispectral data from various sensors have great potential for mapping landscapes, but distinct spectral and spatial resolution capabilities are crucial for accurately classifying wildlife habitats. Our study aimed to develop a technique for precisely discerning habitat categories for the Himalayan Ibex (*Capra sibirica*) using different satellite imagery. To address both spectral and spatial challenges, we utilized LISS IV and Sentinel 2A data and integrated the LISS IV data with Sentinel 2A data along with their corresponding geometric information. Employing multiple supervised classification algorithms, we found the Random Forest (RF) algorithm to outperform others. The integrated (LISS IV-Sentinel 2A) classified image achieved the highest accuracy, with an overall accuracy of 86.17% and a Kappa coefficient of 0.84.

To map the suitable habitat of the Ibex, we conducted ensemble modeling using the Land Cover Land Use (LCLU) of all three image types (LISS IV, Sentinel 2A, Integrated) and other predictors such as topographical, soil type, vegetation, and water radiometric indices. The integrated model provided the most accurate prediction of suitable habitat for the Ibex, surpassing the other two LCLU classes derived from individual images. The Soil Adjusted Vegetation Index (SAVI) and elevation were identified as crucial factors in identifying suitable habitats

These findings hold valuable implications for the development of effective conservation strategies, as accurate classification schemes enable the identification of vital landscape elements. By precisely classifying LULC satellite images and identifying crucial habitats for the Ibex, this pilot study provides a new and valuable strategy for conservation planning. It enhances our ability to preserve and protect the habitat of wildlife species in the mountain ecosystem of the Himalayas.

## 1. Introduction

Satellite imagery is full of detail and contributes significantly to the dissemination of geographical information (Muhammad et al. 2012). The utilisation of satellite and remote sensing images provides both quantitative and qualitative data, which streamlines fieldwork and shortens research times (Chaichoke et al. 2011). Satellite remote sensing techniques capture images at regular intervals. Satellites with various band ranges, geographical resolutions, and spectral resolutions have made it possible to gather more types of remote sensing data from the same area (Wang et al., 2001). The significance of proficient analysis and processing of remote sensing images has been on the rise owing to the exponential growth of remote sensing data (Yin et al. 2021). Because of its wide range of potential applications in fields as diverse as geography, ecology, city planning, forest monitoring, and the military, remote sensing image scene classification is receiving increasing amounts of research and development funding (Cheng et al. 2017). There are several different types of multiresolution and multispectral data now accessible. The data acquired through multispectral remote sensing is characterised by narrow spectral bands that possess a relatively larger bandwidth. Consequently, the gathered data can be employed to examine the spatial characteristics of ground substances (Vohra and Tiwari 2020). The limitations of a single source of satellite data in accurately extracting ground objects are attributed to spectral resemblance among different objects or spatial proximity between the objects. Consequently, for the enhancement of data evaluation precision, it is imperative to appropriately construe object characteristics such as configuration and spatial interconnections, in conjunction with the spectral response (Vohra and Tiwari 2020).

It has become a challenge in remote sensing technology to figure out how to combine different sourced data and produce the most useful image possible. However, in recent years have seen an increase in the importance of image fusion within image processing applications as a result of the plethora of available acquisition methods. Integrating many images from various sensors into a single, useful one for analysis, is the goal of image fusion, which is a relatively a new field (Ma et al. 2019). The comprehension of digital image fusion techniques can enhance the interpretation of multiresolution and multi-sensor data, resulting in improved images that are more suitable for both human perception and seamless computer analysis tasks such as extraction of features, segmentation, and object recognition (Luo et al. 2016; Berger et al. 2015). The fusion of data from multiple sensors and resolutions has been found to be beneficial in enhancing the quality of low-resolution data. Additionally, this approach offers supplementary information collected from the same geographical location, which can be better comprehended than relying solely on data from a single sensor (Ma et al. 2018). Combining the spatial and spectral characteristics of remote sensing images, image fusion technology has gradually broadened its application field, and now the fusion image concept has been applied to land cover land use (hereafter LCLU) categorization. Recently, there has been an explosion of interest in using multi-sensor data fusion for LCLU classification. Accuracy in land cover feature categorization can be improved through the integration of data from many remote sensing sensors with varying resolutions. The demand for greater precision in image and data analysis has spurred research into multiresolution and multi-sensor data, as well as improved methods for gaining access to higher-resolution remote sensing data (Rodriguez-Galiano et al. 2012a). The biophysical state of the Earth’s surface and immediate subsurface is characterised by the composition of topography, soil, surface water, forest, grassland, groundwater, marsh and human structures, collectively referred to as “land cover.” Furthermore, land use for recreational purposes, wildlife habitat, and agricultural land can the example of land use (Turner et al. 1995; Weng 1999; Sherbinin 2002).

It is obvious that land cover classification, derived from remotely sensed data, is still a crucial societal need for natural resource management, surveillance, and development strategies (Colditz et al. 2011; Topaloğlu et al., 2016; Khatami et al., 2016). Numerous studies have generated information from remote sensing by making LCLU maps from various data sources like multispectral, hyperspectral and radar aperture (Craig Dobson et al. 1995; Soria-Ruiz et al., 2010; Pal and Foody 2010; Szuster et al., 2011; Miettinen and Liew, 2011; Srivastava et al., 2012; Hütt et al., 2016; Fonteh et., 2016; Wei et al., 2016; Büyüksalih, 2016; Mohajane et al., 2018; Sirro et al., 2018; Juliev et al., 2019, Camargo et al., 2019, Zafari et al. 2019), subsequently, the LCLU types were classified through the utilisation of machine learning algorithms (Patenaude et al. 2005; Rosenqvist et al. 2003; Wulder et al. 2018). This type of mapping is useful for assessing landuse dynamics, identifying ecosystem services, understanding the effects of global climate change, and formulating land use policy (Fry et al., 2011; Burkhard et al., 2012; Gebhardt et al., 2014; Guidici and Clark, 2017; Noi and Kappas 2017; Hussain et al., 2020).

Furthermore, with the growing statistical models nowadays Species distribution models (hereafter SDM), is a great conservation tool which enhance the capabilities of conservation managers to delaminate conservation priority areas on the base of species presence and the association with their environment. SDM can directs to finds appropriate conservation policies, estimating the area under invasion, evaluate species richness of an area and estimate the probable habitat of any species (Franklin, 2010). SDMs perform crucial role in quantitively ecology by systematically inspecting a species’ relation, in terms of evaluating the biotic and abiotic factors for distributing an organism in a given area (Franklin, 2010). The SDMs approach was developed based on Hutchinson’s ecological niche theory, initially introduced in the 1950s and later revised by Booth et al. (1988). Within the ecological context, the species exhibits significant interactions with various factors such as dietary resources, vegetation types, elevation profiles, and climatic elements (Morris et al., 2012; Besnard et al., 2013). The SDMs employ several methods to determine how a species’ presence in the environment affects that species’ ability to choose an area on a spatial surface that would be favourable for that species (Guisan and Thuiller, 2005; Franklin, 2010; Peterson et al., 2011). It is quite challenging to determine whether a species is actually absent, even though occurrence records can be obtained through museums, published literatures, and field studies. Several SDM methods have been created that solely use positive presence data in order to address this difficulty (Phillips et al., 2006). Instead of using only one modelling technique, the ensemble modelling strategy uses multiple SDM models, which increases the accuracy of predictions about a species’ geographic range (Araújo and New, 2007; Thuiller et al., 2009; Marmion et al., 2009). Due of the ambiguity in selecting one strategy from numerous, ensemble modelling is more effective to a single SDM technique (Pearson et al., 2006; Elith and Graham, 2009; Buisson et al., 2010; Garcia et al., 2011).

Himalayan Ibex, also known as Himalayan Ibex is a member of Bovidae family is a true goat species. This caprinae have a wide distribution range in the mountains of India, Pakistan, China, Mongolia, Afghanistan, Tajikistan, Uzbekistan, Kazakhstan, Kyrgyzstan and Russia (Shackleton 1997, Fox et al.1991, Fedosenko and Blank 2001). This species has a wide residence which ranging from Karakoram and Hindu-Kush mountains of Pakistan through the higher elevated areas of Lahaul-Spiti and some other patches of Himachal Pradesh. The existence of this mountain goat species in this region since 1641 CE as per Saini et al. 2019. This species has very wide range of habitat in its distributional range which mainly configure by steep slopes, mountain ridges, rugged terrain, rocky outcrops, cold deserts and foothills (Dzieciolowski et al. 1980; Clark et al. 2006, Khan et al. 2016). It is the largest among the Capra genus, adult males can reach up to 5.6 feet and weigh up to 130 kg (Fedosenko and Blank 2001). The Himalayan Ibex are affable, and its social groups are composed of both single-sex and mixed-sex individuals, with the latter being observed only during certain seasons. By affecting plant species composition, vegetation structure, and nutrient cycling, ungulates play a significant role in preserving ecosystems (McNaughton 1979; Bagchi and Ritchie 2010). Therefore, maintaining and managing ungulate populations and their habitat is a key goal of conservation management. Habitat is one of the prerequisites components which supports any species to survive. Therefore, monitoring habitat is one of the crucial elements which can help to make any conservation strategy. Himalayan Ibex is an “Near Threatened” mammalian species as per IUCN Red list (Reading et al. 2020), and categories as a Schedule I species under the Wildlife (Protection) Act, 1972 in India. This caprinae species lost their habitat with the rapid urbanization, habitat destruction, hunting and poaching.

The present study aims at the integration of LISS IV and Sentinel 2A images over the Jispa valley of Lahaul Spiti district, Himachal Pradesh, which is under Trans-Himalayan landscape. The purpose of this research is to combine high spatial and spectral data to better distinguish different types of LCLU and find how the classification help to predict the habitat of Himalayan Ibex in this landscape.

## 2. Study area

The present study was carried out in the Jispa valley site. The landmass segment is located in the eastern part of the Lahaul valley in the Lahaul Spiti district of Himachal Pradesh, India (Fig 1). Total area of the study area encompasses 559 km^2^, and lies in UTM zone 43N. This landmass falls under Trans Himalaya Ladakh Mountains (1A) Indian biogeographic zones, which geomorphology is very distinct. High mountains, inclining slopes, and sparse vegetation are the main features of this area. This area has only two clearly defined seasons. High snowfall is frequent during the winter. Farming is one of the primary sources of income in this area; the main commercial crops, which are only grown in the summer, are potatoes, peas, cauliflower, and cabbage. This area is intersected by the River Bhaga. This landmass is extremely important since it supports a variety of crops, biodiversity, pasture, linking roads, and human settlements.

**Figure 1:**
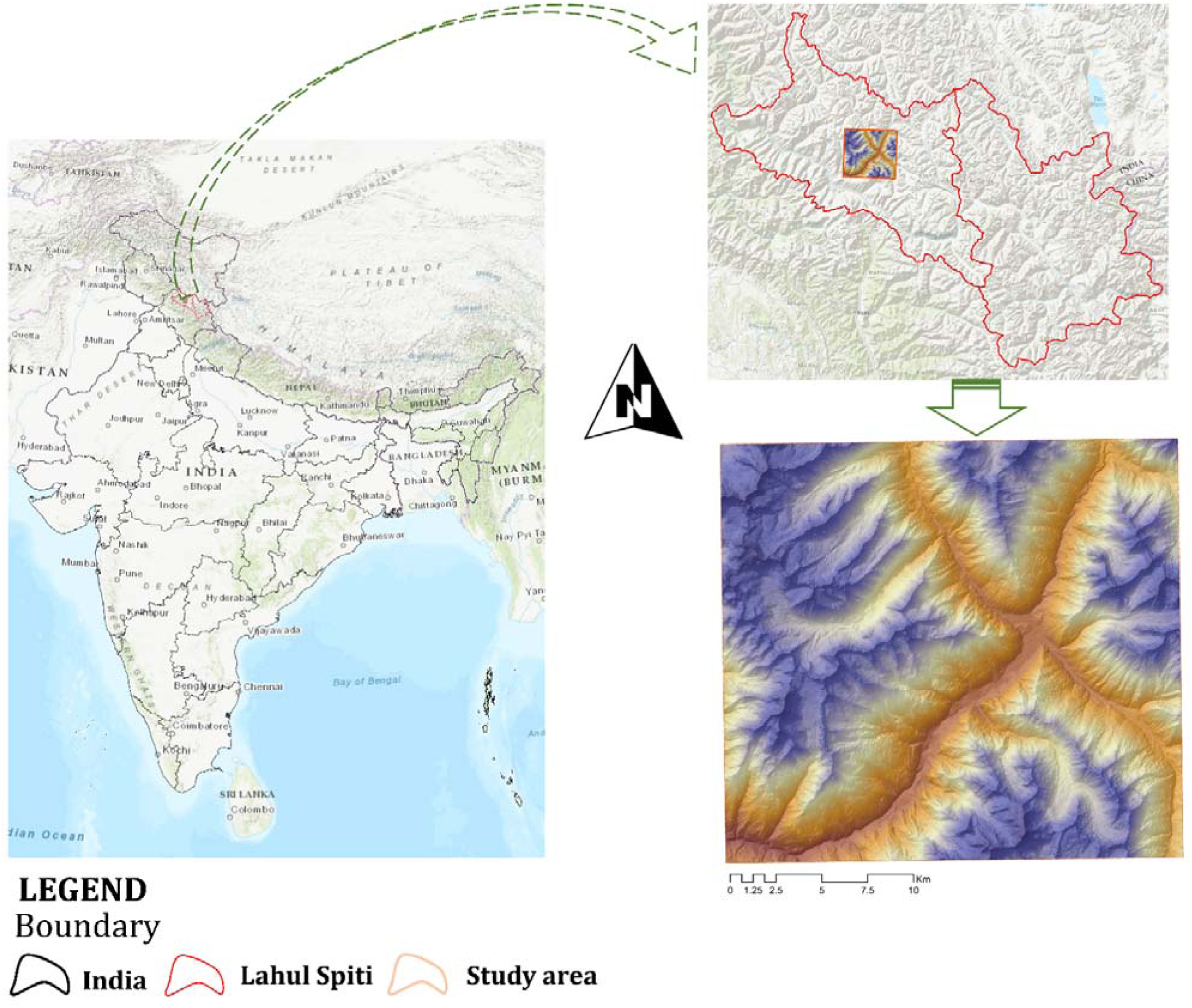
Depiction of present study area along with the its location in India.

## 3. Methodology

### 3.1. Land Class and Land Use Classification using multiple satellite imagery

#### 3.1.1. Data Acquisition

In implementing this study, we assessed two different source satellites viz. Linear Imaging Self-Scanning Sensor (LISS) IV and Sentinel 2A. LISS IV is a multi-spectral sensor with high resolution and data providing in three spectral bands (*viz.* B2 0.52 μm - 0.59 μm, B3 0.62 μm - 0.68 μm, B4 0.77 μm - 0.86 μm). A ground resolution of 5.8 m (at nadir) provides by LISS-IV. Moreover, a rotating deck is attached to a Payload Steering Motor mounted with LISS-IV, which can revolve by 26 degrees and allow for a 5-day revisit of any given ground region. Since the system has 10-bit quantization and can cover 100% of the albedo with a single gain, no gain commands are necessary. Furthermore, Sentinel 2A is an optical multispectral imaging mission with a wide-swath and high resolution. The Global Navigation Satellite System (GNSS), a dual-frequency receiver with an orbital accuracy-specific propulsion system, assists in maintaining each satellite’s position in orbit. The multispectral optical instrument of the Sentinel 2A satellites contains 13 spectral bands (Visible, Near-Infra-Red and Short Wave Infra-Red) with spatial resolutions of 10 m, 20 m, and 60 m for the various spectral bands with a 290 km swath width. The satellite’s sun-synchronous orbit is at a mean altitude of 786 km, and it completes a 5-day cycle with the two satellites (Drusch et al., 2012). LISS IV imagery was collected from Bhoonidhi portal (https://bhoonidhi.nrsc.gov.in/) (Table 1) and Sentinel 2A (Table 1) Level-1C multispectral instrument scenes were downloaded from United States Geological Survey (https://earthexplorer.usgs.gov/).

**Table 1:**
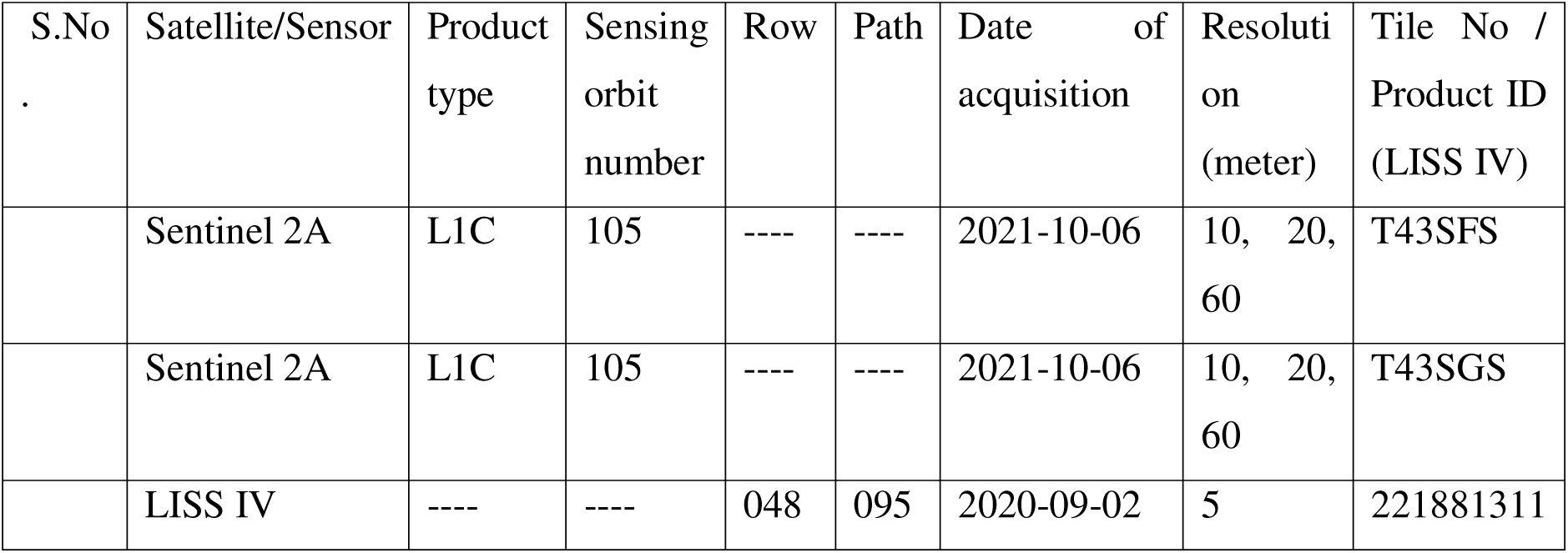
Data type and acquisition details for the two satellite imagery.

The exact geometric correction and registration of two images is the most fundamental requirement for accurate image classification (Balcik, F.B. and Sertel, E., 2002). Sentinel 2A data was pre-processed in the Sentinel Application Platform (SNAP Desktop, Version 6.0.0) for resolution enhancement of the bands by the highest resolution of the other bands and all the bands were converted into the same resolution (Brodu,2017). The satellite image band was stacked in ArcGIS 10.6 environment with layer stacked function. The amalgamation of a higher spatial resolution image and a higher spectral resolution image is advantageous for remote sensing research (Chitade and Katiyar, 2012). Image fusion usually involves two main processes, (a) Image to image geometric registration of two datasets geometry and (b) combining spectral and spatial data to produce a new, enriched dataset that differs from the originals (Lu and Weng, 2007).

Image to image geometric registration is the process of superimposing images (two or more) of the same scene that were captured at various times, from various angles, and or by various sensors (Brown, 1992). The image-to-image geometric registration method was assigned to the satellite images in order to run an accurate fusion procedure. We have used the Coregistration QGIS plugin (Scheffler et al. 2017) for the procedure, where we use the LISS IV image as a reference image and the Sentinel 2A image as a target image.

#### 3.1.2. Integration of the two different satellite images

As we mentioned, we used SNAP for the Sentinel 2A bands to convert the 20 meters bands at the lowest resolution i.e., 10 meters. After that, we applied the nearest neighbour resampling method (Wenbo et al. 2008) to minimize the loss of spectral information of Sentinel 2A images and to gain a spatial resolution of 5 meters, like the LISS IV images. Furthermore, in the integrating image the three bands of LISS IV assembled with the Sentinel 2A bands by the replacement of the 3 bands of Sentinel 2A (i.e., B3, B4, B8; Green, Red and NIR bands respectively). Since, the LISS IV has B2, B3 and B4 (Green, Red and NIR, respectively) bands and the B2, B3 and B8 of Sentinel 2A bands have a similar spectral resolution. After the integration of the LISS IV and Sentinel 2A bands total bands were 10, now it has good spatial and spectral resolution as well as it gains bands number.

#### 3.1.3. Image Classification

The extraction and interpretation of valuable information from enormous satellite images necessitate an effective and efficient statistical methodology. Image classification is a method that involves the categorization of each pixel in an image or raw information obtained from remote sensing satellites, with the aim of generating an appropriate set of labels for specific land cover themes (Lilles and, Keifer 1994; Abburu and Golla 2015; Karlsson 2003).

In supervised satellite image classification methods, the training sample is the most crucial element. Analyst input is required for supervised classification methods. The fundamental process of supervised classification involves the examination of input data, generation of training sets and signature files, and the assessment of the training sets and signature files’ efficacy (Abburu and Golla 2015).

In this study we classify the images (LISS IV, Sentinel 2A, and the integrated image) by five supervised machine learning classification techniques viz. Maximum Likelihood, K-nearest neighbourhood, Support Vector machine, Guassian mixed model and Random Forest algorithm. In this present study the images classified into nine classes of LCLU types: agriculture land, sparse vegetation, barren land, scrub areas, juniper patch, settlements, permafrost, water bodies and road ways. The training data utilised in this classification process was gathered for all eight LCLU classes, with the exception of permafrost, through field surveys conducted within the study landscape. We have used the same training polygons to classify the LISS IV, Sentinel 2A as well as the integrated image.

The idea is to evaluate the performance of various satellite images using their bands as variables and to calculate the precision of mapping quantification by applying remote sensing data to actual ground-truth circumstances. The accuracy evaluation of each defined class is determined by an error matrix that compares map information with reference data and the sampled area or points. Such errors are attributed to the accuracy of the producer and user (Congalton and Green 2019; Foody 2002). The accuracy is derived from a final classification error matrix made up of many multivariate statistical studies, where the overall accuracy, kappa, and F-statistics represent the accuracy of various classified classes (Congalton et al., 1983; Congalton 1991; Foody 2002; Nitze et al. 2012; Keshtkar et al. 2017, Gumma et al. 2019).

As a parametric statistical and supervised classification method, the Maximum Likelihood (hereafter ML) classifier is the most popular technique used in remote sensing applications (Jia et al. 2011). This approach uses an average with values, variance, and covariance classification technique that is based on statistics and takes into account the values of the variables (Günlü 2021). However, Gaussian mixture model (hereafter GMM) is another machine learning algorithm. They are employed to divide data into various groups in accordance with the probability distribution. GMMs uses to classify satellite images based on probabilistic concept (Gu et al. 2007; Chellappa et al. 2009; Okwuashi et al., 2011; Lakshmi et al. 2015). One of the simplest machine learning and supervised learning methods is the K-nearest neighbour algorithm (hereafter KNN). The KNN algorithm is a method for classifying objects by utilising the proximity of training samples in the feature space (Bremner et al., 2005, Nurwauziyah et al., 2018). Support Vector Machine (hereafter SVM) is essentially a supervised machine learning technique. The structural risk minimization concept and statistical learning theory serve as the foundation for the SVM (Günlü 2021). This algorithm is typically used to address classification-related challenges, while it can also be used to address regression-related complications. SVM differentiates between the objects based on the separation of the hyper-planes (Huang and Zhang,2012, Luo et al., 2015, Melgani and Bruzzone 2004).

It has been proved that Random Forest (hereafter RF) is capable of producing accurate LCLU maps (Rodriguez-Galiano et al., 2012b; Sonobe et al., 2014). RF constructs decision trees before randomly combining them (Breiman 2001; Liaw & Wiener, 2002; Ishwaran & Kogalur, 2007; Ishwaran et al., 2008; Noi and Kappas 2017, Sonobe et al., 2017). The supervised classification of these three images performed by dzetsaka classification tool, SCP tool in QGIS and ArcGIS 10.6 (Esri 2018, Karasiak 2019, Congedo 2021)

### 3.2. Habitat Suitability modelling for Ibex

#### 3.2.1. Occurrence locations of the Himalayan Ibex

During the extensive field study during 2019-2022 we recorded the occurrence of Himalayan Ibex. The collection of the occurrences of the species’ we followed camera trapping method, trail sampling, vintage point and questionnaire survey. We have sampled all types of habitats in the search of Himalayan Ibex. However, the study area composed of rugged terrain, steep slopes, high mountains, uncertain weather conditions so representative sampling was carried out. During the field work we gathered 167 presence locations, furthermore we used spatially autocorrelated 82 locations for final analysis.

#### 3.2.2. Variables preparation and selection

The variable selection for SDM is the critical stage in this analysis because the variables should be plausible for the Himalayan Ibex presence. The Trans-Himalayan landscape is mostly rugged and found gradients in elevation with less amount of vegetation growth which makes this area complex and this complexity also observe in the habitat selection by wild animals like Himalayan Ibex. We used Digital elevation model data from Alos Palsar at 12.5- meter spatial resolution and used as primary source for derived slope and aspect. The LCLU classes were generated from the good accuracy produced by classified images. Furthermore, radiometric indices of soil, vegetation and water computed from Sentinel 2A image. All the data was rasterized and resampled at same spatial scale i.e. at 5 meter by the help of spatial analyst extension tool of ArcGIS 10.6. Primarily we prepared 23 ecological pertinent variables for this study, however, for the final model building only selected uncorrelated variables through Pearson correlation coefficients (r) higher than 0.8.

#### 3.2.3. Selection of performing models and evaluation of modeling algorithms

Because of the complex and varied nature of the association between species and their environmental predictors, studies indicate that no single modelling approach is best in every circumstance (Marmion et al., 2009). Modelling algorithms broadly classified as Classification models, Regression models and Complex models. Hence, we applied Multivariate adaptive regression splines (MARS) and Generalized linear model (GLM) from Regression models, Boosted regression trees (BRT) from Classification models and Maximum Entropy Model (MaxEnt), Random Forest (RF) from Complex models, each form with 10-fold cross-validation (Hayes et al. 2015, Dutta et. al 2022). We developed the modelling workflow using the SAHM module by utilising the VisTrails pipeline; using this method, models were able to choose the predictors that best describe the model performance (Morisette et al., 2013; Talbert and Talbert, 2012, Dutta et al. 2022). Each model generated an estimate of the potential habitat suitability for every pixel, presented as continuous values ranging from 0 to 1, which was subsequently interpreted to represent the probable habitat suitability for a specific pixel in the current study area. The predicted habitat suitability was determined through the binary maps, with consideration given to the minimal training occurrence as a threshold (Hayes et al., 2015). We generated 3 ensembled maps as follows, LISS IV derived LCLU and other topographic and radiometric variables at 5-meter spatial resolution, Sentinel 2A derived LCLU and other topographic and radiometric variables at 10 meter spatial resolution, Integrated image derived LCLU and other topographic and radiometric variables at 5 meter spatial resolution. The average of all of the binary predictions (0 or 1) of the five models are used in the ensemble maps for each map pixel, moreover the count surface depicts the model agreement, where 0 refers no model agreement and 5 refers all model agreement for suitability estimation (Dutta et al. 2022).

For the purpose of comparing the performance of SDMs, numerous performance metrics are frequently utilised. We employ area under the receiver operating characteristic curve (AUC), Cohen’s Kappa, True Skill Statistic (TSS), Proportion Correctly Classified (PCC), sensitivity, and specificity to more clearly grasp comparative SDM performance across the 5 model forms (Cohen, 1968; Allouche et al., 2006; Phillips 2010; Illán et al. 2010; Jiménez-Valverde et al. 2013). Generally, AUC considered as a threshold-independent statistic to evaluate models (Guisan and Zimmermann 2000; Brotons et al. 2004; Elith et al. 2006; Phillips et al. 2006; Pearson 2007; Grenouillet et al. 2011). We used the minimal training presence threshold for metrics (specificity and sensitivity), which are threshold-dependent (Peterson et al. 2011). The criterion for building ensemble model was based on AUC threshold value of >0.75 of CV dataset. Variable importance was evaluated by the mean AUC, which is the AUC values ratio and calculated by number of models runs for every model.

## 4. Result

### 4.1. Classification of Land Class and Land Use

The outcome of the three classified images (LISS IV, Sentinel 2A and the integrated image) quality assessment by different accuracy assessments have been discussed in the following sections. The improvements in the integrated image have been examined by comparing the integrated products with the LISS IV and Sentinel 2A images. The integrated image has a 5 metre spatial resolution like LISS IV, but it also improved spectral signature. We derived nine LCLU classes of the study area from the three types of images with five different supervised classification algorithms (Fig 2). 810 sample points (90 points for each class) across the study area were obtained by the stratified random sampling approach to evaluate the classification accuracy of the images. The study region consists of several LCLU features such as sparse vegetation, scrub covers, juniper patches, road lines, water, agriculture field, human settlements, barren land, permafrost area. Many of the aforementioned properties are challenging to extract from LISS IV sensor data because of its fine resolution, which fail to distinguish many classes at once. With the aid of integrated data, the classification result increases and the features identified more accurately. As a result, fused data might be a cost-effective alternative to better resolution multispectral data. The classification accuracy of integrated classified images was compared to that of LISS IV and Sentinel 2A classified images. Classification accuracy estimated by overall accuracy (OA), Kappa coefficients (κ) Weighted Kappa, Producer’s accuracy (PA), User’s accuracy (UA), commission, omission and F-statistics (F) (Table 2, 3, S1 – S4). (Congalton 1991). Result depicts that the classification accuracy not good by LISS IV image, no classifying algorithm perform well, the highest overall accuracy achieved by SVM (OA = 60.61) and κ statistic calculated was 0.56, and the lowest performance by GMM algorithm (OA= 48.76). and κ statistic calculated was 0.42, so the two classifying algorithms achieve moderate accuracy (Table 2, Fig S1 -S5) (Landis and Koch 1977). Moreover, the other classier viz. ML, RF and GMM classifier overall accuracy score was 57.40, 56.41, 48.76 and κ statistic score was 0.52, 0.51, 0.42 respectively that directed also moderate accuracy on the LISS IV image (Table 2, Fig S1 - S5).

**Figure 2:**
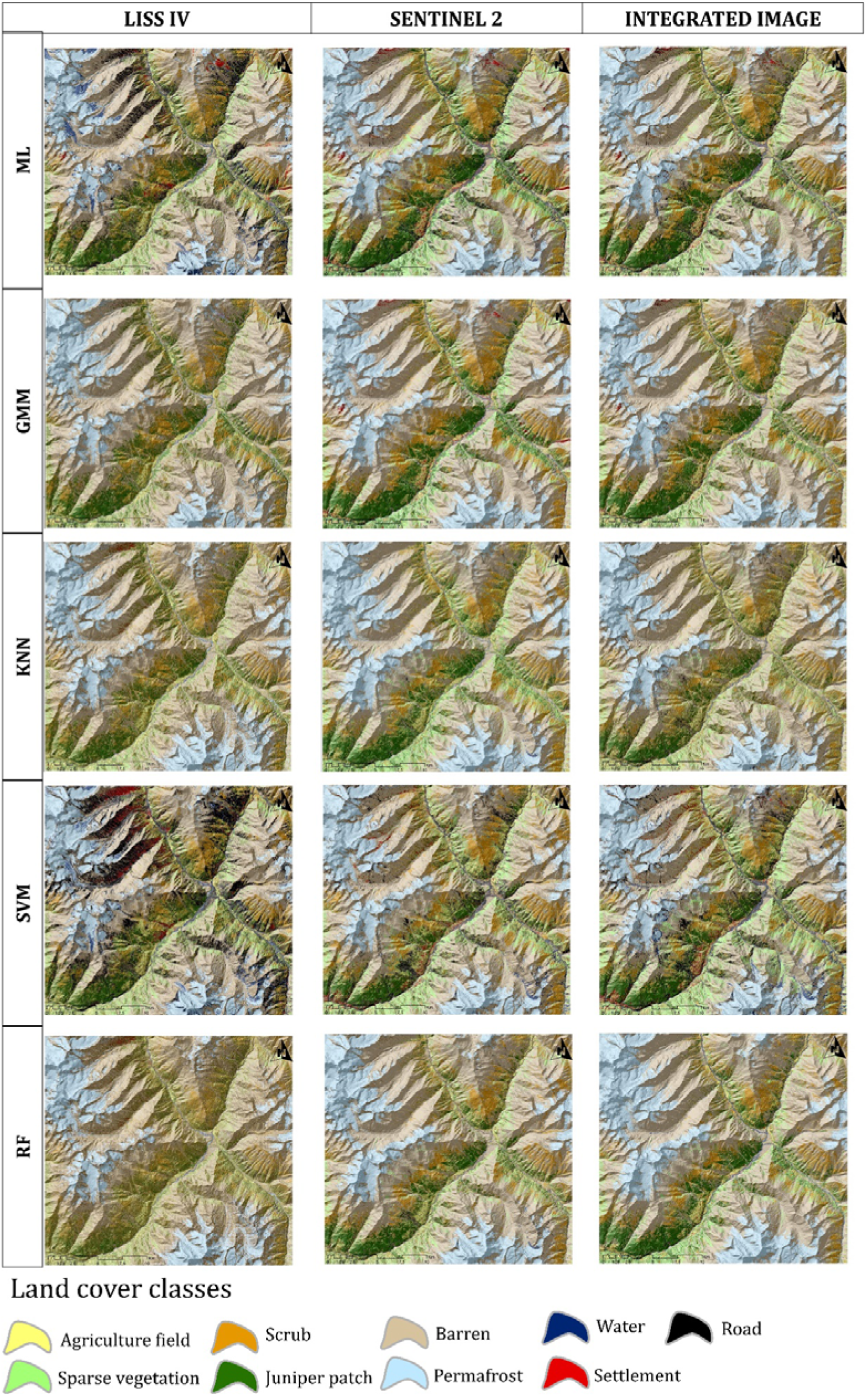
Classified images from LISS IV, Sentinel 2A and Integrated images by ML, GMM, KNN, SVM and RF classifier algorithms.

**Table 2:** Accuracy assessment of LISS IV, Sentinel 2A and Integrated images classified by five supervised classifiers namely, ML (Maximum Likelihood), GMM (Guassian mixed model), KNN (K-nearest neighbourhood), SVM (Support Vector machine) and RF (Random Forest). The evaluation metrices are Overall accuracy, Kappa (κ) statistic, SE of Kappa (κ) statistic and Weighted Kappa (κ).

| Accuracy Assessment | Image type | ML | GMM | KNN | SVM | RF |
| --- | --- | --- | --- | --- | --- | --- |
| Overall accuracy | LISS IV | 57.41 | 48.77 | 55.31 | 60.62 | 56.42 |
|  | Sentinel 2A | 73.08 | 71.35 | 70.24 | 76.54 | 80.24 |
|  | Integrated | 71.73 | 67.78 | 81.85 | 76.79 | 86.17 |
| Kappa ($\kappa$) statistic | LISS IV | 0.52 | 0.42 | 0.50 | 0.56 | 0.51 |
|  | Sentinel 2A | 0.70 | 0.68 | 0.67 | 0.74 | 0.78 |
|  | Integrated | 0.68 | 0.64 | 0.80 | 0.74 | 0.84 |
| SE of Kappa ($\kappa$) statistic | LISS IV | 0.02 | 0.02 | 0.02 | 0.02 | 0.02 |
|  | Sentinel 2A | 0.02 | 0.02 | 0.02 | 0.02 | 0.02 |
|  | Integrated | 0.02 | 0.02 | 0.02 | 0.02 | 0.01 |
| Weighted Kappa ($\kappa$) | LISS IV | 0.51 | 0.41 | 0.46 | 0.57 | 0.50 |
|  | Sentinel 2A | 0.71 | 0.65 | 0.60 | 0.72 | 0.74 |
|  | Integrated | 0.71 | 0.61 | 0.80 | 0.74 | 0.85 |

The accuracy estimation of the classification algorithm on Sentinel 2A imagery evaluate that the RF model classify the area with highest overall accuracy i.e., 80.24 and κ statistic 0.78 and the lowest performance by KNN algorithm i.e., 70.24 and κ statistic 0.67, so the results show substantial agreement (Table 2, Fig 2, Fig S1 -S5). Other algorithms viz. SVM, ML and GMM performing OA score 76.54, 73.08 and 71.35 respectively and the κ statistic score was 0.73, 0.70, 0.68, respectively which is also substantial accurate as per Landis and Koch (1977) (Table 2, Fig 2, Fig S1 -S5). Interestingly, the integrated image shows higher classification accuracy compared to LISS IV and Sentinel 2A image (Table 2, Fig 2, Fig S1 -S5). The highest overall accuracy produces by RF i.e., 86.17 and κ statistic 0.84 which is a perfect accuracy estimation (Table 2, Fig 2, Fig S1 -S5). KNN classifier also perform great with the integrated image classification than the LISS IV and Sentinel 2A with an overall accuracy value of 81.85 and κ statistic 0.80 and again it qualifies as perfect classification (Table 2, Fig 2, Fig S1 -S5). The other classification algorithm such as the SVM, ML and GMM predicted overall accuracy are 76.79, 71.72, 67.77 respectively and κ statistic 0.74, 0.68, 0.64 respectively, which qualify as substantial classify (Table 2, Fig 2, Fig S1 -S5). From this overall accuracy and κ statistic it is clear that integrated image classification performs well than the single sensor images. However, the weighted κ also depicts the same (Table 2, Fig S1 -S5).

However, the accuracy estimation for identifying each feature class from these images we calculated producer’s accuracy, user’s accuracy F- statistics, omission and commission error rates. Regarding to the comparison of each feature class, permafrost class differentiate by each classification algorithms on every image perfectly. Omission error rates depicts that the highest pixels misclassified by GMM algorithm on LISS IV image (Table 3, Table S1 -S4, Fig 2, Fig S1 – S5). Road, settlement and sparse vegetation classes are poorly classified by this algorithm on the LISS IV image (Table 3, Table S1 -S4, Fig 2, Fig S1 – S5). However, the commission error rate directs that the road, highly misclassified by ML classification on the LISS IV image (Table 3, Table S1 -S4, Fig 2, Fig S1 – S5). However, classification of LISS IV image by ML, GMM, SVM poorly classify settlement, scrub, juniper patch, barren and agriculture classes (Table 3, Table S1 -S4, Fig 2, Fig S1 – S5). The result of producer’s accuracy shows that road, settlement and sparse vegetation, agriculture and water poor accuracy in terms of classified by GMM algorithm on LISS IV image (Table 3, Table S1 -S4, Fig 2, Fig S1 – S5). The user’s accuracy and F- statistics depicts that the road, settlement and sparse vegetation, agriculture and water class gained highest accuracy with the integrated image classified by RF algorithm (Table 3, Table S1 -S4, Fig 2, Fig S1 – S5). Furthermore, the KNN classifier on integrated image and RF classifier on Sentinel 2A image having overall accuracy score >= 80%, κ statistic and weighted κ score >=0.80, which tells perfect object classification but the best classification by RF on the integrated image among all the classification (Table 3, Table S1 -S4, Fig 2, Fig S1 – S5).

**Table 3:** Class accuracy metrices on best classified image i.e., Random Forest classifies the Integrated image. The class accuracy metrices evaluated by Omission rate, Commission rate, Producer’s accuracy, User’s accuracy and F – stat.

|  | LCLU Class | LISS-IV | Sentinel 2A | Integrated |  | LCLU Class | LISS-IV | Sentinel 2A | Integrated |
| --- | --- | --- | --- | --- | --- | --- | --- | --- | --- |
| Omission | Agriculture | 47 | 23 | 14 | Commission | Agriculture | 47 | 27 | 16 |
|  | Sparse vegetation | 57 | 26 | 20 |  | Sparse vegetation | 40 | 17 | 11 |
|  | Barren | 13 | 2 | 1 |  | Barren | 55 | 38 | 33 |
|  | Scrub | 42 | 7 | 18 |  | Scrub | 57 | 20 | 22 |
|  | Juniper patch | 36 | 17 | 13 |  | Juniper patch | 59 | 33 | 17 |
|  | Settlement | 70 | 34 | 23 |  | Settlement | 16 | 3 | 4 |
|  | Permafrost | 2 | 0 | 0 |  | Permafrost | 21 | 1 | 0 |
|  | Water | 34 | 11 | 3 |  | Water | 9 | 1 | 1 |
|  | Road | 91 | 58 | 31 |  | Road | 33 | 14 | 5 |
|  | LCLU Class | LISS-IV | Sentinel 2A | Integrated |  | LCLU Class | LISS-IV | Sentinel 2A | Integrated |
| Producer's Accuracy (PA) | Agriculture | 53 | 77 | 86 | User's accuracy (UA) | Agriculture | 53 | 73 | 84 |
|  | Sparse vegetation | 43 | 74 | 80 |  | Sparse vegetation | 60 | 83 | 89 |
|  | Barren | 87 | 98 | 99 |  | Barren | 45 | 62 | 67 |
|  | Scrub | 58 | 93 | 82 |  | Scrub | 43 | 80 | 78 |
|  | Juniper patch | 64 | 83 | 87 |  | Juniper patch | 41 | 67 | 83 |
|  | Settlement | 30 | 66 | 77 |  | Settlement | 84 | 97 | 96 |
|  | Permafrost | 98 | 100 | 100 |  | Permafrost | 79 | 99 | 100 |
|  | Water | 66 | 89 | 97 |  | Water | 91 | 99 | 99 |
|  | Road | 9 | 42 | 69 |  | Road | 67 | 86 | 95 |
|  | LCLU Class | LISS-IV | Sentinel 2A | Integrated |  |  |  |  |  |
| F- statistics | Agriculture | 53 | 75 | 85 |  |  |  |  |  |
|  | Sparse vegetation | 50 | 78 | 84 |  |  |  |  |  |
|  | Barren | 60 | 76 | 80 |  |  |  |  |  |
|  | Scrub | 49 | 86 | 80 |  |  |  |  |  |
|  | Juniper patch | 50 | 74 | 85 |  |  |  |  |  |
|  | Settlement | 44 | 78 | 85 |  |  |  |  |  |
|  | Permafrost | 88 | 99 | 100 |  |  |  |  |  |
|  | Water | 76 | 94 | 98 |  |  |  |  |  |
|  | Road | 16 | 57 | 80 |  |  |  |  |  |

### 4.2. Habit Suitability Ensemble model

After applying supervised classification on these satellite imageries and evaluate the accuracy of image classification we used these LCLU to evaluate the potential habitat distribution of Himalayan Ibex, in this study landscape. Habitat suitability of a species can address by its environment where the species thrive. The LCLU one of the important aspects which tells about the habitat of the species. Performing the species distribution model is crucial for conservation of species in their natural habitat. Our model builds with 82 uncorrelated occurrence records of Himalayan Ibex. Furthermore, we assigned the LCLU derived from the best classified image of three different images (Sentinel 2A, LISS IV and Integrated image). Only LCLU are not enough when the species resides in complex and rough terrain. So, we choose topographical variables and soil, vegetation and water radiometric indices along with the LCLU for knowing the suitable habitat of this species. Classified LCLU maps helps to understand the habitat more precisely, fine resolution can provide more insights to the habitat of the species. However, we selected 21 predictors at starting (Table S5), after the multicollinearity test, we removed correlated variables and use only variables which follow the Pearson’s rule i.e., <0.8 (Fig S6 – S8). The AUC values of five different modelling algorithm ranged from 0.77-0.92 when habitat class use from LISS IV classified image, AUC values ranged from 0.77-0.91 when habitat class use from Sentinel 2A classified image and when integrated classified image derived habitat class use for the suitability prediction the AUC value ranged from 0.77-0.92 for training data sets, which reflects great performance by the models (Table 4, Fig 3). The model performance also evaluated by other metrices like True skill statistic (TSS), Percent Correctly Classified (PCC), Cohen’s Kappa, specificity and sensitivity and depicts good performance by the models (Table 4, Fig S9 – S14). The development of the final ensemble model and ensemble count maps was achieved as every participation model satisfied the AUC requirement of 0.75 and above (Fig 3, 4). The LISS IV derived LCLU with other variables show most suitable area about 63.80 km^2^, the Sentinel 2A derived LCLU with other variables show most suitable area about 72.42 km^2^ and the Integrated image derived LCLU with other variables show increase in most suitable area about 78.42 km^2^ (Table S6, Fig 6, 7). In all instances when the rest of the model uses some of the variables, RF and MaxEnt methods used all uncorrelated variables. Furthermore, for all of the models with all of the combinations, the Soil Adjusted Vegetation Index (SAVI) with the highest mean AUC, found as a major contributor and most significant variable among the variables (Fig S15 – S32). Different LCLU shows significance differently in the three types of images (Fig S15 – S32). In the LISS IV image the Barren is the leading variable which influence in the model building (Fig S15, S18-22), moreover Juniper patch shows higher significance among the LCLU while using the Sentinel 2A and Integrated image (Fig S16 - 17, S23-32). When the Himalayan Ibex occurrence is more prevalent, the response curve for these variables has high peaks, and the likelihood values fall as the distance increases (Fig 18 - 32).

**Figure 3:**
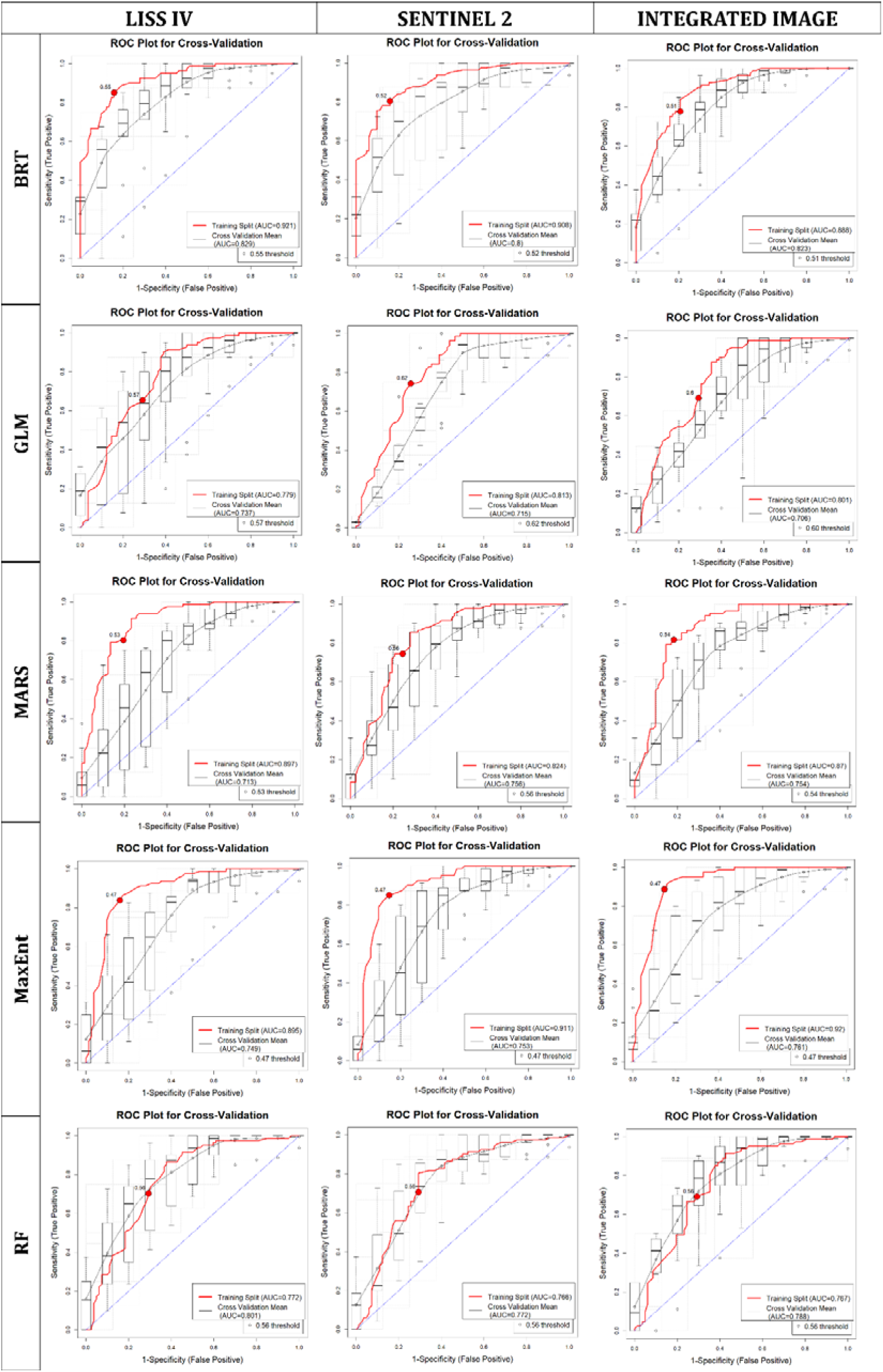
AUC plots of five algorithms on different sourced images to predict species distribution model.

**Figure 4:**
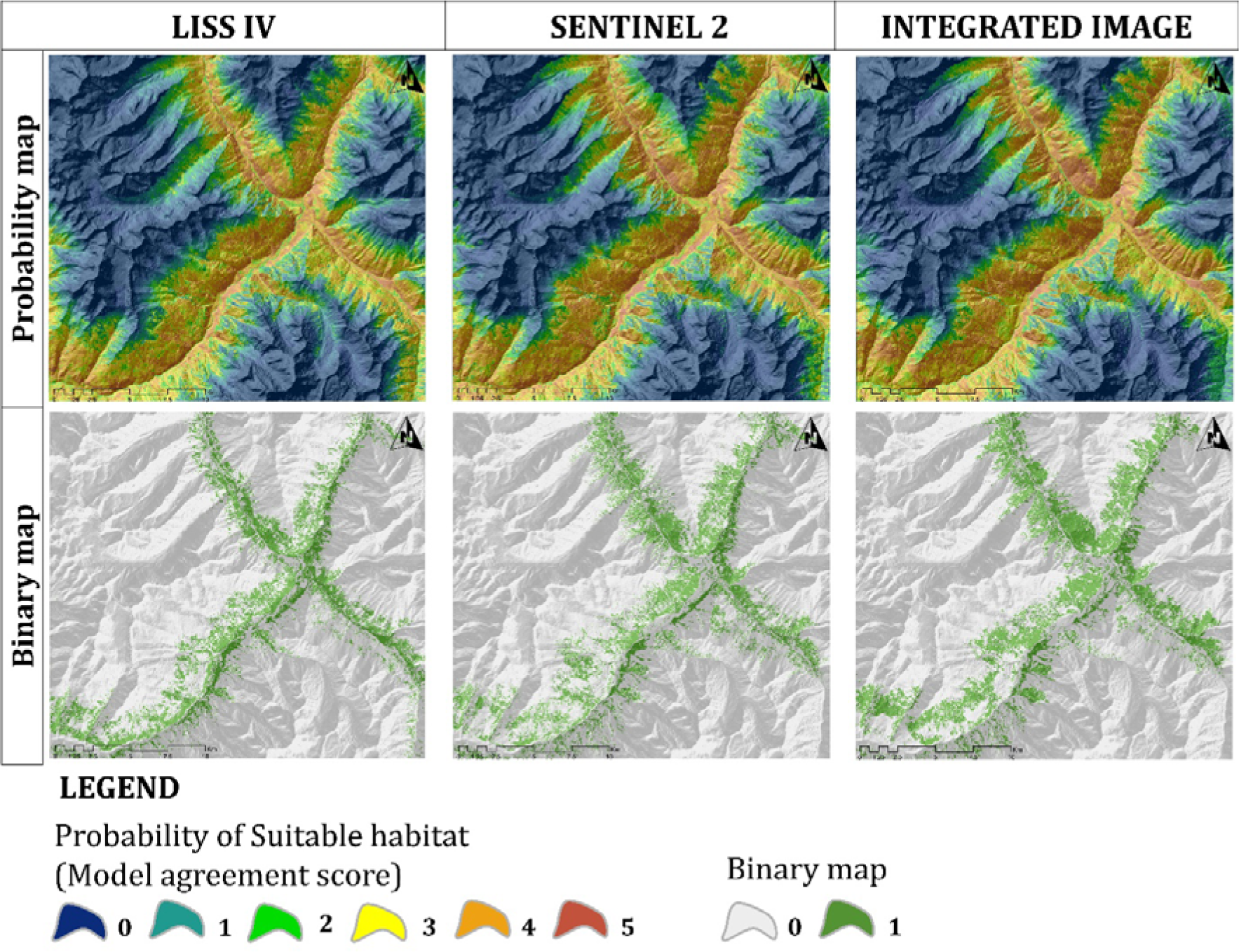
Probability (model agreement maps) maps and binary maps from three different sourced images generated LCLU to predict species distribution model of Himalayan Ibex.

**Table 4:** Evaluation metrics to evaluate the efficiency of the participating distribution models for Himalayan Ibex in the study landscape. Participating models are BRT (Boosted Regression Tree), GLM (Generalized Linear Model), MARS (Multivariate adaptive regression splines), MAXENT (Maximum Entropy Model), RF (Random Forest) and the efficiency of the models evaluated by AUC (area under the receiver operator curve), PCC (Proportion Correctly Classified), sensitivity, specificity, Cohen’s kappa and TSS (True Skill Statistic). CV mean (Cross Validation) data used for model evaluation and train split used for the model, which was assessed by model evaluation. (a) LISS IV derived LCLU used for this model building, (b) Sentinel 2A derived LCLU used for this model building and (c) Integrated image derived LCLU used for this model building.

| Model | AUC | PCC | Sensitivity | Specificity | Kappa | TSS |
| --- | --- | --- | --- | --- | --- | --- |
| BRT | 0.92 | 84.67 | 0.85 | 0.84 | 0.69 | 0.69 |
| GLM | 0.78 | 68.1 | 0.65 | 0.71 | 0.36 | 0.36 |
| MARS | 0.9 | 80.37 | 0.8 | 0.8 | 0.61 | 0.61 |
| MAXENT | 0.9 | 83.95 | 0.84 | 0.84 | 0.68 | 0.68 |
| RF | 0.77 | 70.55 | 0.7 | 0.71 | 0.41 | 0.41 |

(b)
| Model | AUC | PCC | Sensitivity | Specificity | Kappa | TSS |
| --- | --- | --- | --- | --- | --- | --- |
| BRT | 0.91 | 82.32 | 0.8 | 0.84 | 0.65 | 0.65 |
| GLM | 0.81 | 74.39 | 0.74 | 0.74 | 0.49 | 0.49 |
| MARS | 0.82 | 75 | 0.74 | 0.76 | 0.5 | 0.5 |
| MAXENT | 0.91 | 85.28 | 0.85 | 0.85 | 0.71 | 0.71 |
| RF | 0.77 | 70.73 | 0.71 | 0.71 | 0.41 | 0.41 |

(c)
| Model | AUC | PCC | Sensitivity | Specificity | Kappa | TSS |
| --- | --- | --- | --- | --- | --- | --- |
| BRT | 0.89 | 78.53 | 0.78 | 0.79 | 0.57 | 0.57 |
| GLM | 0.8 | 69.94 | 0.69 | 0.71 | 0.4 | 0.4 |
| MARS | 0.87 | 81.6 | 0.81 | 0.82 | 0.63 | 0.63 |
| MAXENT | 0.92 | 87.04 | 0.89 | 0.85 | 0.74 | 0.74 |
| RF | 0.77 | 69.94 | 0.69 | 0.71 | 0.4 | 0.4 |

## 5. Discussion

The theme of this research was to compare the object classification methods derived from different sensor imagery and the preparation of an integrated image and analysis the classification performance of it, for the improvement of LCLU analysis which play pivotal role for species habitat configuration. Two different sensors namely LISS IV and Sentinel 2A, were considered for the preparation of the integration image. The images were classified by 5 different type of supervised classification technique viz. ML, GMM, KNN, SVM and RF. As a result, five LISS IV, five Sentinel 2A and five integrated images were produced (Fig 2). In total fifteen images were compared with each other by qualitatively measures (Table 2, 3, S1-S4, Fig S1-S5). Consequently, all of the images were subjected to visual interpretation and comparison. Subsequently, the integrated images revealed discrete outcomes that evince enhancements of varying magnitudes (Table 2, 3, S1-S4, Fig S1-S5). The results of the qualitative evaluation indicate that RF produced the most advantageous result on integrated images, while GMM produced the least favourable results on LISS IV images (Table 2).

Classified images by integration techniques were investigated also by comparing the classification results of original LISS IV and Sentinel 2A images for each selected class (Table 2, 3, S1-S4, Fig S1-S5). The results demonstrate a clear improvement in classification when utilising the RF method on the integrated image. However, KNN, SVM and ML of Integrated showed great accuracy estimation for the image classification. Moreover, classifier algorithms perform also well in the Sentinel 2A image after integrated image (Table 2, 3, S1- S4, Fig S1-S5). In the case of LISS IV image, the classifier did not perform good, which easily interpret by the accuracy estimation. Among the all models GMM classification on LISS IV image was the worst (Table 2, 3, S1-S4, Fig S1-S5). In a nutshell it has been determined that the utilisation of multi-sensor data significantly improves the precision of LCLU classification, resulting in a more dependable and superior map generation. The integration of two different satellite image result showed promising result to differentiate the LCLU types. All the supervised classification methods used to classify the images discussed in this paper are capable of properly classify multispectral images with good accuracy except LISS IV. Furthermore, some model imperfectly classify settlement, road, water bodies which account by visual interpretation (Fig 2).

The habitat suitability model of Himalaya Ibex, derived from the best classification method of three images depicts there is a change in most suitable habitat (Table S6, Fig 4-5). From this study we conclude that in this rugged terrain the species have complex association with habitat which can not only defined by land classes, it also highly influenced by the physical character of the terrain (Fig S15-S32). From the distribution model assessment, we found that SAVI and elevation have major role which shape their habitat (Fig S15-S32). Area with little vegetation cover, SAVI is used to adjust the NDVI for the impact of soil brightness. So, the improperly classified image of LISS IV predict least suitable area followed by Sentinel 2A classified image (Table S6, Fig 4-5). Furthermore, the integrated image derived LCLU with the other topographic and radiometric variables predict an increase in suitable area (Table S6, Fig 4-5). SDM with all image derived LCLU and other variables predicts 34.76 km^2^ common (Table S6, Fig 4-5). Whereas, 41.50 km^2^ area common in LISS IV and Sentinel 2A derived LCLU SDM model, 44.91 km^2^ area common in LISS IV and integrated image derived LCLU SDM model and 54.14 km^2^ area common in Sentinel 2A and integrated image derived LCLU SDM model (Table S6, Fig 4-5). Therefore, the study’s findings show that an integrated image can distinguish LCLU more effectively than a single satellite image of the two images employed in this study. The present study evaluates the performance of the SDM in conjunction with the three image-derived LCLU datasets, incorporating both topographical and radiometric variables. The results indicate that the SDM exhibits favourable performance across all LCLU datasets. Notably, the integrated image dataset demonstrates a high level of precision in identifying LCLU, which may prove useful in characterising the habitat configuration of the Himalayan Ibex.

**Figure 5:**
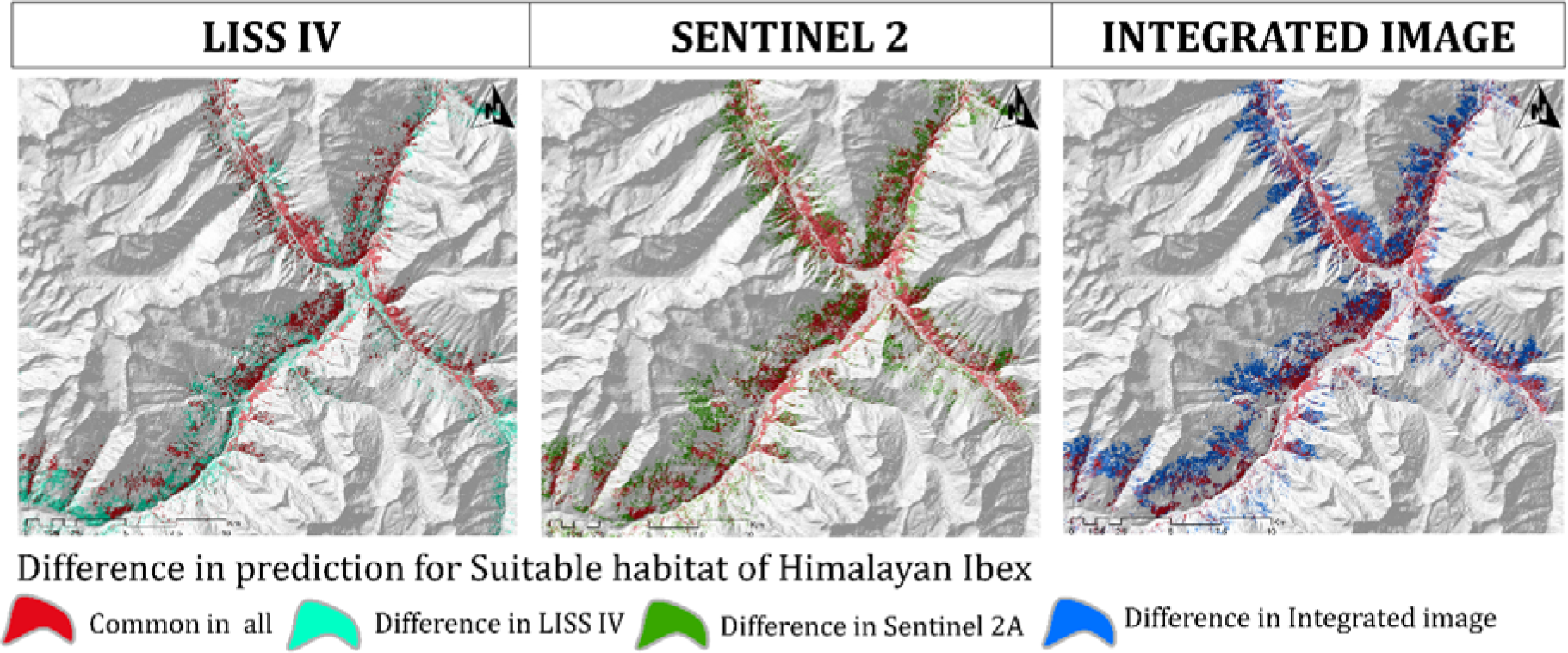
Variance in suitable habitat prediction of Himalayan Ibex by three different sourced images generated LCLU.

### 5.1. Conclusion

The current study deal with the comparison between the contribution of LISS IV, Sentinel 2A, and the integrated image of these aforementioned multisensor image with five classification algorithms for the enhancement of LCLU analyses. After that, we select the best classified image of these three different types of images for extraction of LCLU and predict suitable habitat of Himalayan Ibex and compare which is the best among them.

For the image classification we employed five supervised classification algorithms, namely ML, GMM, KNN, SVM and RF. Subsequently, statistical analysis and visual interpretation were performed on each image. Due to the heterogeneous landscape’s complexity, the classes can occasionally be difficult to distinguish; nevertheless, the integrated image classified by RF method significantly enhanced the mapping of each class. The F- statistics, Producers accuracy, user’s accuracy, commission and omission rate show that the classes classify with low accuracy scores in LISS IV and Sentinel 2A images, however, employing integration to improve their accuracy. In conclusion, the integrated image showed unique findings with varying degrees of improvement. For both visual and qualitative analysis of the classified images, RF demonstrated the best results on integrated image and GMM classification on LISS IV was the worst. From the accuracy assessment the second-best classifier algorithm was KNN on integrated image. However, it is evident that Sentinel 2A image also perform good to classify the ground objects but no classifier algorithm could not classify LISS IV image at a significant way.

Furthermore, this investigation reinforces the efficacy of SDM ensemble modelling in the prioritisation of conservation strategies and management. We recommended ensemble model, against using a single modelling technique, particularly for species with complex habitat. Due to the geomorphological complexities of this study area, only LCLU is not sufficient to predict this species suitable habitat, so the topographic and radiometric variables useful for this study. However, it can conclude that how the specific topographic factor with LCLU supports the habitat of the species potential distribution in this study area, because the landcover classes distributed in specific altitudinal gradients. The maximum models use elevation and SAVI for prediction of suitable habitats in this area. However, we can infer from this study that topographical characteristics may serve as a proxy for identifying suitable habitat of mountainous species when LCLU classification is not fair, hence the poorly classified image of LISS IV also shows fair suitable areas of the species.

Himalayan Ibex is a mountain ungulate which help to shape the ecosystem by many dimensions. The snow leopard, the apex predator in this Trans Himalayan environment, relies on this species of mountain ungulate as one of its primary food sources. (Suryawanshi et al. 2017, Sharief et al. 2022). Furthermore, the indigenous inhabitants of this region have several kinds of traditional beliefs regarding Himalayan Ibex.

We therefore draw the conclusion from this study that all types of images can be helpful for species distribution models in this type of complicated terrain when they are associated with other crucial and influential variables like elevation, SAVI, slope, and aspect. However, a precise description of LCLU can help to know the distribution of suitable habitat in space more significantly, which helps to understand where the focus for conservation needs to be placed for the long-term survival of wild animals. With this finding, we suggested a community conservation area in the study landscape because it is clear that there is a considerable amount of habitat for this wild mountain goats in this area, despite the fact that there is no such protected area. It can also be advantageous for the inhabitants and the natural population of Himalayan Ibex to safeguard this area from unforeseen infrastructures because it can offer various ecosystem services, such as water, sustainable tourism, and opportunities for outdoor recreation.

## Notes

### Competing Interest Statement

The authors have declared no competing interest.

